# Telomere-to-telomere *Schizosaccharomyces japonicus* genome assembly reveals hitherto unknown genome features

**DOI:** 10.1101/2023.09.04.556195

**Authors:** Graham J Etherington, Pei-Shang Wu, Snezhana Oliferenko, Frank Uhlmann, Conrad A. Nieduszynski

## Abstract

*Schizosaccharomyces japonicus* belongs to the single-genus class Schizosaccharomycetes, otherwise known as ‘fission yeasts’. As part of a composite model system with its widely studied *S. pombe* sister species, *S. japonicus* has provided critical insights into the workings and the evolution of cell biological mechanisms. Furthermore, its divergent biology makes *S. japonicus* a valuable model organism in its own right. However, the currently available short-read genome assembly contains gaps and has been unable to resolve centromeres and other repeat-rich chromosomal regions. Here we present a telomere-to-telomere long-read genome assembly of the *S japonicus* genome. This includes the three megabase-length chromosomes, with centromeres hundreds of kilobases long, rich in 5S ribosomal RNAs, transfer RNAs, long terminal repeats, and short repeats. We identify a gene-sparse region on chromosome 2 that resembles a 331 kb centromeric duplication. We revise the genome size of *S. japonicus* to at least 16.6 Mb and possibly up to 18.12 Mb, at least 30% larger than previous estimates. Our whole genome assembly will support the growing *S. japonicus* research community and facilitate research in new directions, including centromere and DNA repeat evolution, and yeast comparative genomics.

**Take-away:**

- A telomere-to-telomere genome assembly of the fission yeast *S. japonicus*
- Chromosome 2 harbours a previously unknown second centromere-like region
- The estimated genome size of *S. japonicus* may be up to 18.12 Mb

## Introduction

Fission yeasts are a single-genus group of yeasts within the Taphrinomycotina subdivision of Ascomycota fungi, which divide equatorially into two daughter cells of equal size (Taylor & Berbee, 2006). The *Schizosaccharomyces* genus contains a well-established model organism *S. pombe*, together with *S. octosporus, S. cryophilus, S. osmophilus, S. lindneri, S. versatilis*, and an emerging model system *S. japonicus*. All known species of fission yeast have now been sequenced (Brysch-Herzberg et al., 2023; Jia et al., 2023; Rhind et al., 2011; Wood et al., 2002).

*S. pombe* is a long-established model organism utilised across a wide range of cellular and molecular biology research (Rutherford, Harris, Oliferenko, & Wood, 2022; Wood et al., 2002; Wood et al., 2012). Despite being closely related and relying on many conserved genes, several major differences with *S. japonicus* has made this pair of sister species emerge as a powerful evolutionary cell biology system (Alam, Gu, Reichert, Bahler, & Oliferenko, 2023 ; Gu, Yam, & Oliferenko, 2015; Makarova et al., 2016; Makarova et al., 2020; Yam, He, Zhang, Chiam, & Oliferenko, 2011). Furthermore, *S. japonicus* has become a standalone model organism for study of processes not present or tractable in other yeasts (Aoki et al., 2011; Chapman, Taglini, & Bayne, 2022; Furuya & Niki, 2010, 2012; Gomez-Gil et al., 2019; Gu & Oliferenko, 2019; Kinnaer, Dudin, & Martin, 2019; Lee et al., 2020; Nozaki, Furuya, & Niki, 2018; Papp, Acs-Szabo, Batta, & Miklos, 2021; Pieper, Sprenger, Teis, & Oliferenko, 2020; Rutherford et al., 2022; Wang et al., 2021; Yam, Gu, & Oliferenko, 2013).

A chromosome-level assembly of *S. japonicus* totalling 11.7 Mb has long been available (Rhind et al., 2011) and has recently been utilised to create a valuable online database and research tool for this species – JaponicusDB (Rutherford et al., 2022). The assembly of the three *S. japonicus* chromosomes was inferred from genetic linkage within the seven largest supercontigs which were then joined with four stretches of N’s. Three of these joins are centromeres, which were not resolved in the assembly and were substituted by stretches of ‘N’s, 120 kb in length. The final gap, on chromosome 2 between supercontigs 7 and 5, was the last region to be assembled (Rhind et al., 2011). This join lies roughly 400 kb downstream from the centromere; it contains four stretches of ‘N’s between 100 bps and 1,680 bps in length.

The availability of low-cost long-read sequencing combined with advances in genome assembly algorithms has allowed for chromosome-scale telomere-to-telomere sequencing to become achievable even over complex genomes (Belser et al., 2021; Nurk et al., 2022; Tong et al., 2019). Here we utilise Oxford Nanopore (ONT) long-read sequencing to assemble a telomere-to-telomere reference sequence of *S. japonicus* and compare it to the current version of the reference genome (GCF_000149845.2_SJ5).

## Materials and Methods

### Cell culture and high molecular weight DNA extraction

High molecular weight genomic DNA was isolated from a mating type h-isolate of *S. japonicus* strain ATCC10660 (Furuya & Niki, 2009), using the Nanobind CBB DNA extraction kit (PacBio), according to the manufacturer’s instructions with modifications. Briefly, *S. japonicus* was cultured in YE media supplemented with 3% glucose, adenine hydrochloride (100 mg/L), histidine (75 mg/L), leucine (75 mg/L) and uracil (75 mg/L) at 30 °C. Forty millilitre culture of exponentially growing cells at OD_600_ = 0.5 were harvested, washed twice with 0.05M EDTA pH 8.0 and once with CSE buffer (50 mM citrate-phosphate pH 5.6, 40 mM EDTA and 1.2 M sorbitol). The cells were then treated with zymolyase 100T (0.6 mg/ml in CSE buffer) at 37 °C for 50 minutes. The subsequent cell lysis and DNA extraction were performed as described in the manufacturer’s protocol for hypocrealean fungi, except that CLE3 buffer was replaced with an optimized Proteinase K buffer (10 mM Tris-HCl pH 9.5, 0.5 M EDTA, 125 mM Na_2_SO_3_, 1% lauryl sarcosine), while Buffer SB was replaced with 10 mM Tris-HCl, pH 9.5. The size distribution of the extracted genomic DNA was confirmed by pulsed field gel electrophoresis.

### DNA sequencing

DNA concentration and size were assessed using the double-stranded high-sensitivity assay on a Qubit fluorometer and a TapeStation system using a Genomic DNA ScreenTape, respectively. Genomic DNA was sequenced using the GridION nanopore sequencer (Oxford Nanopore Technologies). One sequencing library was generated using a ligation kit (SQK-LSK112) and sequenced on one MinION R10.4 flow cell (FLO-MIN112), following the manufacturer’s guidelines. This generated 3.38 Gb of QC-passed ONT long read data (from 498,295 reads), with an estimated N50 read length of 27.31 kb. 646 reads were greater than 100kb in length, 9 reads were greater than 200kb, and the maximum read-length was 284 kb.

### Additional genomic datasets

We used four libraries of publicly available *S. japonicus* Illumina short-read data for sequence polishing (BioProject accession PRJNA770288). We also downloaded the current publicly available genome assemblies and annotations for *S. japonicus* from NCBI (GCF_000149845.2_SJ5), and JaponicusDB (Rutherford et al., 2022). Although originating from the same initial reference sequence, the two different data sources contain complementary data. The NCBI annotation contains information such as gene biotypes and products, whilst the JaponicusDB reference sequence is assembled into named chromosomes (1-3) and the mitochondrial genome.

### Genome assembly

We used Alvis (Martin & Leggett, 2021) to align ONT reads to the *S. japonicus* reference assembly. This identified 4982 reads that appeared to be chimeric, which were removed from the dataset. Then, we used Canu (Koren et al., 2017) to correct and trim the ONT reads and then generated three *de novo* assemblies; one using all the data and one each using reads >50 kb and >60 kb. We then assessed the three assemblies on their completeness and contiguity, expecting the ‘best’ assembly to be that which resolved each of the three *S. japonicus* chromosomes with the fewest contigs as well as assembling the mitochondrial genome. We removed contigs from the assembly that Canu suggested were graph bubbles (‘suggestBubble=yes’ in the Canu fasta header) and trimmed contigs that were suggested to be circular (‘suggestCircular=yes’ in the Canu fasta header). Further, we performed a self-versus-self blastn of the mitochondrial genome and trimmed any redundant overlapping sequence (Altschul, Gish, Miller, Myers, & Lipman, 1990). Finally, we polished the final assembly with the four libraries of Illumina data using three rounds of Pilon (Walker et al., 2014).

Next, we used the nucmer tool in the MUMmer tool suite (Delcher, Salzberg, & Phillippy, 2003) to align contigs in the assembly to that of the *S. japonicus* genome from JaponicusDB to identify syntenic chromosomal regions between the two assemblies, visualising the output in Dot (Nattestad, 2020).

We carried out quality control steps to assess the final assembly. As well as calculating contig-specific GC content and contiguity statistics, we also used BUSCO (v5.3.2) (Manni, Berkeley, Seppey, & Zdobnov, 2021) to identify the number of Ascomycota-specific single-copy orthologs and KAT (Mapleson, Garcia Accinelli, Kettleborough, Wright, & Clavijo, 2016) to compare the distribution of k-mers in the high-accuracy Illumina reads (from BioProject accession PRJNA770288) to that of the assembly. To compare repeat content between our assembly and that of JaponicusDB, we ran a repeat masking pipeline with RepeatModeler v2.0.3 (Flynn et al., 2020) and RepeatMasker (v4.1.2-p1) (Smit, Hubley, & Green, 2015; Tarailo-Graovac & Chen, 2009) and summarised the results using the ParseRM.pl script from the Parsing-RepeatMasker-Outputs tool (https://github.com/4ureliek/Parsing-RepeatMasker-Outputs).

### Genome annotation

We annotated coding and non-coding genes as follows. Liftoff (v1.6.3) (Shumate & Salzberg, 2020) was used to lift over the NCBI *S. japonicus* annotation (GCF_000149845.2_SJ5) to our new assembly, allowing extra gene copies with a CDS sequence identity of 95% and over to be annotated. Using the output from RepeatMasker (see above), the generated GFF file was divided into several repeat-specific categories, namely short, A-rich, GA-rich, and LTRs (including *ab initio* predictions). Additionally, we calculated the concentration of these repeats in defined regions of the assembly (e.g., centromeres, telomeres, unplaced contigs, etc).

tRNAscan-SE (v2.0.12) (Chan, Lin, Mak, & Lowe, 2021) was used to identify tRNAs *de novo*, and barrnap (v0.9) (Seeman, 2018) and rnammer (v1.2) (Lagesen et al., 2007) were used to identify rRNA genes *de novo*.

To identify small nuclear RNA (snRNA) and small nucleolar RNA (snoRNA) genes, we extracted the co-ordinates for all genes of each type from the NCBI *S. japonicus* annotation and for each type used BEDTools getfasta (v2.30.0) (Quinlan, 2014) to extract the sequences. After filtering out sequences greater than 300 nucleotides in length, we used MUSCLE (v3.8.31) (Edgar, 2004) to align the sequences and created a consensus sequence from the alignment using Mega11 (Tamura, Stecher, & Kumar, 2021). Using RNAalifold (Bernhart, Hofacker, Will, Gruber, & Stadler, 2008) we input the consensus sequence to generate a consensus secondary structure for the alignment. Next, we used Infernal (v 1.1.4) (Nawrocki & Eddy, 2013) to reformat the aligned sequences into Stockholm format and manually added the consensus secondary structure as the metadata entry for ‘#=GC SS_cons’. Using the Stockholm formatted alignment file, we used Infernal cmbuild and cmcalibrate to build and calibrate our snRNA and snoRNA models and used each model to search our assembly for snRNAs and snoRNAs *de novo*, reformatting the output into GFF format. We repeated this pipeline using snRNAs and snoRNAs from the *S. pombe* genome and using ‘BEDTools intersect’, merged the output from the pipeline with that from the *S. japonicus* output, to produce unique non-overlapping snRNA and snoRNA loci. The generated annotation files from all these tools can be found in the supplementary data (Supplementary data S1)

### Comparison to previous S. japonicus assemblies

#### Misassemblies

We used REAPR (v1.0.18) (Hunt et al., 2013) to map paired-end Illumina reads from *S. japonicus* (SRR16290165) to assess assembly errors across both our assembly and that of the JaponicusDB assembly. Using the Smalt aligner, REAPR maps each set of paired-end reads independently to a reference sequence and then breaks assemblies over regions it considers as being misassembled.

#### Whole genome alignments

We used nucmer (Delcher et al., 2003) to align the previously allocated syntenic chromosomes to those in JaponicusDB and examined large breaks in the assembly to identify regions where our assembly had resolved more complex regions of the genome than JaponicusDB. Particular attention was paid to telomeres, centromeres, and other large areas of sequence not present in the JaponicusDB assembly.

#### Telomeres

To calculate the amount of extra telomeric sequence generated, we used ‘delta-filter’ from the MUMmer tool-suite to filter the nucmer alignments for the longest consistent set of alignments and then ‘show-coords’ to provide a human readable version. We then took the alignments, reformatted them into BED format and used BEDTools merge to merge non-overlapping alignments that were within 100bps of each other. Then, using the aligned co-ordinates, we calculated the theoretical 5’ and 3’ extension of the telomeres (compared to JaponicusDB), taking into account JaponicusDB sequences that were not part of the alignment. For example, if an alignment between our assembly and JaponicusDB ended at position 5000, but the JaponicusDB sequence had an extra 2 kb of unaligned sequence to the 3’ end of the alignment, we extended the alignment length outwards to position 7000 and calculated the length of unaligned sequence projecting beyond the 3’ tip of the JaponicusDB telomere. We also performed this for the 5’ end of the alignments, extending the alignments to position 1 of the JaponicusDB sequence and calculating the length of unaligned sequence projecting beyond the 5’ tip of the JaponicusDB telomere.

#### Centromeres

The inferred centromeres in JaponicusDB were linked by stretches of ‘N’s, 120 kb in length. To identify the amount of additional centromeric sequence in our assembly, we identified the closest protein-coding genes (with an allocated gene standard name) flanking the centromeres in JaponicusDB, calculated the amount of genomic sequence (including N’s) between the end and start of the two genes and then compared that to the distance between the same genes in our assembly. Note that here we use the term ‘centromere’ to describe a region between the chromosome arms that is sparse in, or devoid of, protein-coding genes (and often rich in repeat content).

#### Coverage

For regions of interest (e.g., centromeres, rRNA arrays, large insertions, etc), we calculated long-read coverage across the regions using the following pipeline. We used minimap2 to map the ONT reads to our assembly using the pre-set parameters for mapping ONT genomic reads, and then Samtools ‘view’ (Danecek et al., 2021) to remove alignments marked as secondary. Next, we used Samtools ‘depth’ to count per-base coverage across all chromosomes, coding-regions of each chromosome arm (defined by the start of the first and end of the last BUSCO single-copy ortholog hit for each chromosome arm), and that of any other given interval of interest and then calculated the mean per-base coverage and standard deviation (SD) across the interval.

## Results

### A telomere-to-telomere S. japonicus genome assembly

Using ONT reads >50 kb, we obtained an assembly that contained three chromosome-sized contigs and one circular contig representing the mitochondrial genome. In total, the assembly comprises of 17 contigs, with an assembly size of 16,601,825 bases (after Pilon polishing) and a contig N50 of 5.27Mb. Then, we aligned our assembly to the JaponicusDB assembly and identified three contigs that each spanned the whole length of one JaponicusDB chromosome and the mitochondrial genome. On this basis we renamed each contig in the assembly to ‘Chr1’, ‘Chr2’, ‘Chr3’, and ‘Mt’ (Figure 1). Finally, we took the mitochondrial genome and trimmed off redundant overlapping ends. We refer to this final assembly as *S. japonicus* ‘EI 1.0’.

**Figure 1.**
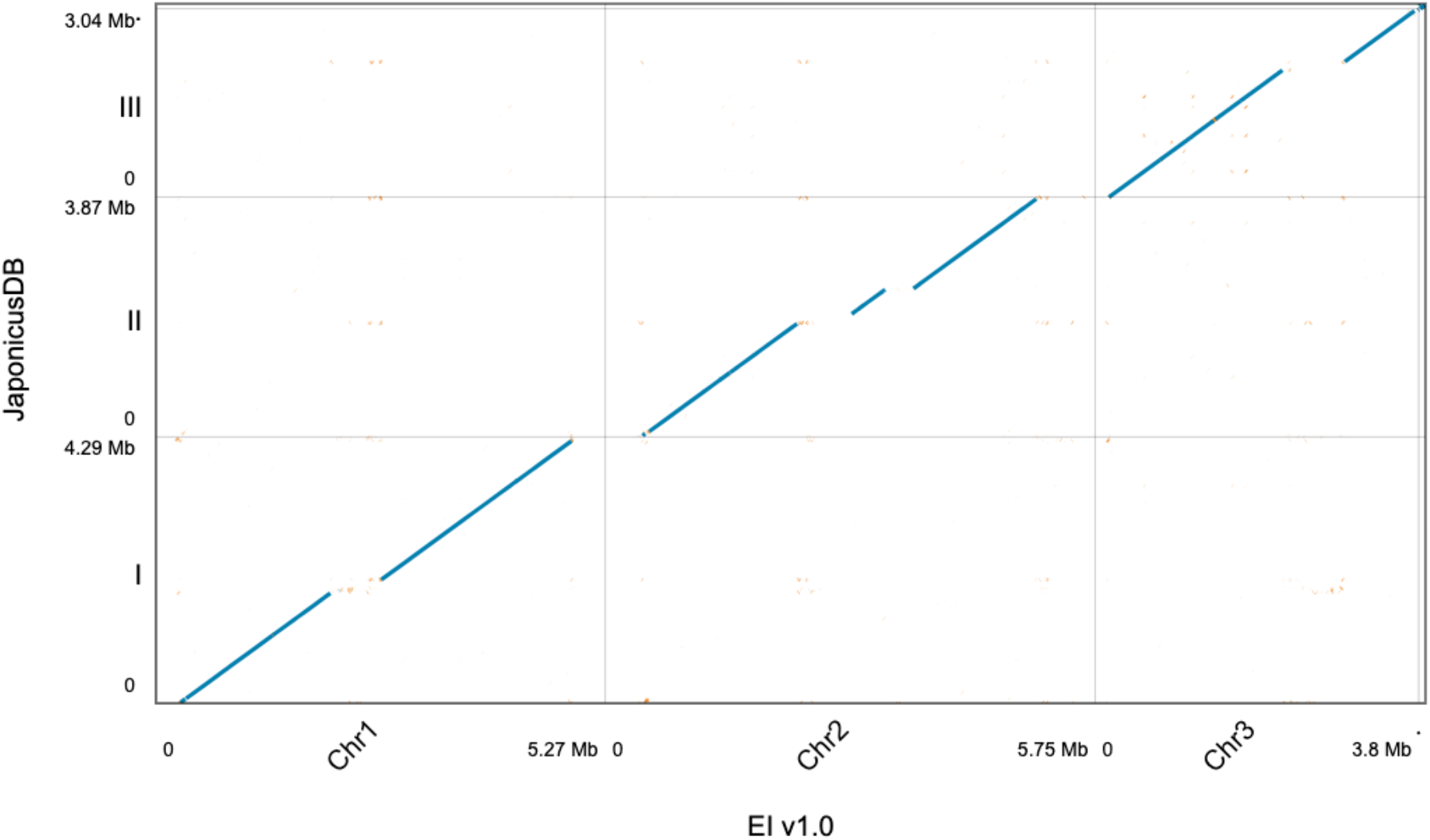
Alignment of the JaponicusDB assembly (y-axis) to EI 1.0 (x-axis). In order to align JaponicusDB assembly with EI 1.0, JaponicusDB was broken at the centromeres, with an additional break on Chr2. JaponicusDB assembly has been scaffolded with N’s to join supercontigs across these areas.

### Distribution of k-mers shows low level of error

We compared the k-mers in *S. japonicus* Illumina short-reads to k-mers present in the EI 1.0 assembly. The distribution of k-mers in the reads found only once in the assembly shows a normal distribution across the graph (Figure 2 in red). As is usual, there are high counts of low-copy k-mers present in the reads but not in the assembly (the black distribution to the left of Figure 2). These k-mers are likely sequencing errors in the Illumina reads that are not reflected in the EI 1.0 assembly. The black distribution of these k-mers extends along the x-axis of the graph. This represent ever decreasing k-mers present in in the reads, but not present in the assembly and likely represents a small number of errors in the ONT reads not corrected by Pilon polishing.

**Figure 2.**
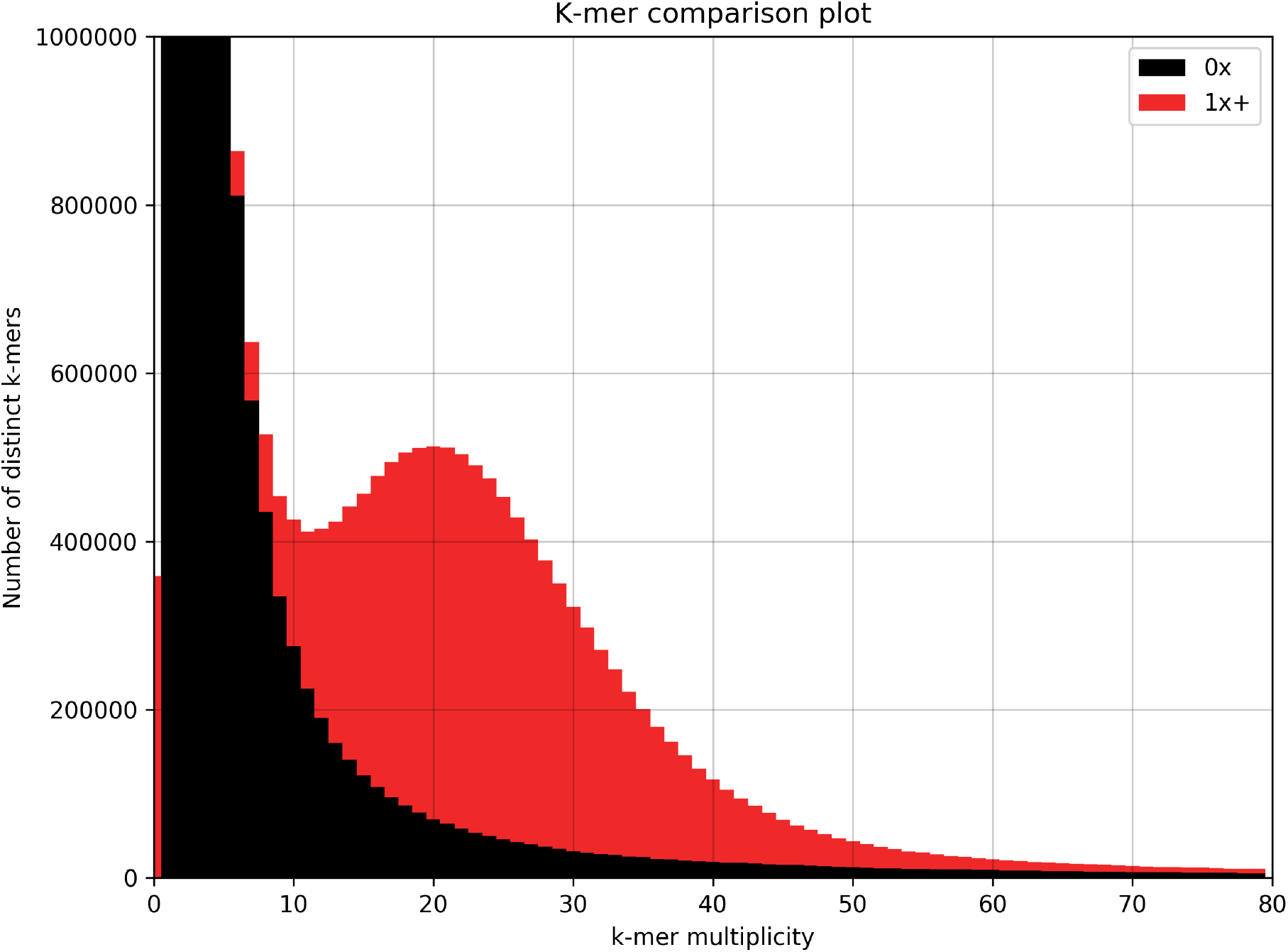
Distribution of Illumina short-read k-mers present once (red) and not present (black) in the EI 1.0 *S. japonicus* assembly. The black peak to the left of the figure represents k-mers in the Illumina reads present in small numbers that have not been incorporated into the final assembly, and probably represent read sequencing errors.

### Composition and chromosome association of unplaced contigs

In addition to the three chromosomes and mitochondrial genome, we retained 13 unplaced contigs that were not marked as assembly bubbles. These additional contigs ranged between 71.3 kb and 240.8 kb in length and had a combined length of 1.7 Mb (Table 1).

**Table 1.**
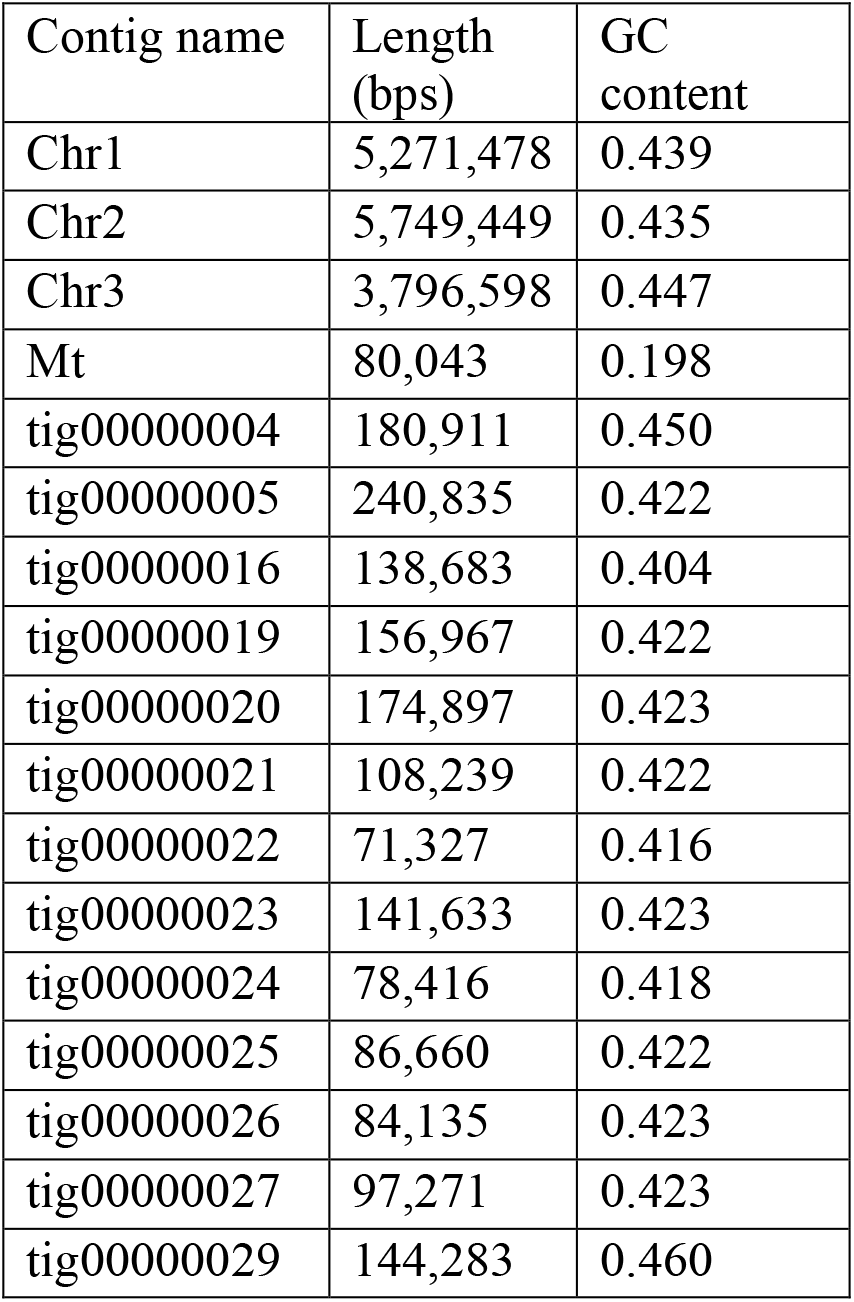
Contig-specific lengths and GC-content for each sequence in the *S. japonicus* EI 1.0 assembly. The mean GC content of the 16 non-mitochondrial contigs was 0.428 and 0.44 across the three chromosomes.

tig00000005 has protein coding genes SJAG_05105, SJAG_05106, and SJAG_06608. SJAG_05105 and SJAG_05106 are both present in one copy and located in JaponicusDB on the unplaced supercont5.6. SJAG_05105 also has an orthologous gene in *S. pombe* (SPAC212.06c), which is located in the telomere of chromosome I. Therefore, this contig might be an unplaced telomeric sequence from Chr 1.

tig00000004 and tig00000029 have one and three copies of protein coding gene SJAG_06608 respectively, which is also present in 19 copies across the 3 chromosomes, and one copy on tig00000005. These two contigs also contain large tRNA arrays.

tig00000022 contains coding genes SJAG_04834, SJAG_04835, and SJAG_04836 at the far 3’ end, along with an 18S-28S rRNA array stretching from the start of the 5’ end towards the coding genes. These three coding genes, along with the same pattern of rRNAs can be found at the end of the short-arm of Chr 3 on both EI 1.0 and JaponicusDB, so this contig appears to be a reverse complement assembly duplication from the short-arm of chromosome 3.

tig00000019, tig00000020, tig00000021, tig00000023, tig00000024, tig00000025, tig00000026, and tig00000027 all contain 18S-28S rRNA arrays across the whole length and are probably all part of the telomeric rRNA array from the short-arm of chromosome 3.

tig00000016 contains no genes or rRNA arrays but is GTCTTA-rich across the whole contig. This sequence matches the previously reported telomeric repeat for *S. japonicus* suggesting that this contig might be unplaced telomeric repeat sequences (Rhind et al., 2011).

### EI 1.0 shows a similar resolution of single-copy orthologs to JaponicusDB

We ran BUSCO to examine the number of single-copy orthologs recovered from both EI 1.0 and JaponicusDB. The results are quite similar, with a small increase (0.7%) of complete and single-copy orthologs recovered from EI 1.0 compared to that of the JaponicusDB assembly (Table 2). This suggests that both assemblies are equally complete in their gene content, and that very little of the assembly content is present in two or more copies.

**Table 2.** Comparison of the number of single-copy orthologs identified in EI 1.0 and JaponicusDB. ‘Complete and duplicated’ orthologs are an indication of duplication in the assembly (e.g., through incorrectly assembling a region of the genome two times or more), ‘fragmented’ refer to genes that are identified but not full-length, and ‘missing’ is an inferred metric of those genes not found in the assembly.

| BUSCO category | EI 1.0 (%) | JaponicusDB (%) |
| --- | --- | --- |
| Complete | 1360 (79.7) | 1349 (79.0) |
| Complete and single-copy | 1308 (76.7) | 1297 (76.0) |
| Complete and duplicated | 52 (3.0) | 52 (3.0) |
| Fragmented | 45 (2.6) | 49 (2.9) |
| Missing | 301 (17.7) | 308 (18.1) |
| Total | 1706 | 1706 |

### Resolution of repeat content accounts for much of assembly size increase

We ran a repeat masking pipeline using RepeatModeler and RepeatMasker and summarised the results (Table 3 and Table 4). We compared the repeat content of JaponicusDB to EI 1.0 (Table 3). EI 1.0 has almost five times more resolved repeat content, comprising almost 15% of the genomic content of which 11.56% are attributable to LTRs (predominantly Gypsy-type). By far the biggest difference between the two assemblies is the Helitron-like transposons that are 29 times more numerous in the EI 1.0 assembly, followed by LTRs, being 5.2 times more numerous.

**Table 3.** Comparison of assembly repeat content (number of nucleotides masked and % of the genome) by class between JaponicusDB and EI 1.0. Abbreviations: SINE -Short interspersed nuclear elements; LTR -long terminal repeat; RC – rolling circle transposons (Helitron-like transposons).

| Repeat class | JaponicusDB | EI 1.0 |
| --- | --- | --- |
| SINE | 20709 (0.18%) | 17450 (0.10%) |
| LTR | 369855 (3.13%) | 1919713 (11.56%) |
| RC (Helitron) | 11497 (0.09%) | 334938 (2.02%) |
| Unknown | 101426 (0.86%) | 200590 (1.21%) |
| **Total masked** | **503,487 (4.26%)** | **2,472,691 (14.89%)** |

**Table 4.** Counts of repeats identified by the RepeatModeler -RepeatMasker pipeline.

| Repeat type | Count |
| --- | --- |
| Short | 4479 |
| LTR | 4362 |
| A-rich | 600 |
| G-rich | 246 |
| GA-rich | 95 |

The pipeline identified 9782 different repeats (where the repeat consisted of at least two bases), of which short repeats were the most numerous (4479). 482 of the short repeats were present in a single tandem array. The top three repeats present in the greatest number of copies (‘CTAAGA’, ‘TTAGTC’, and ‘GTCTTA’) all represent 797 repeat copies varying around one repeat (and its reverse complement) – GACTAA, previously reported as the *S. japonicus* telomeric repeat (Rhind et al., 2011).

Additionally, we calculated the span of repeats in defined regions of the assembly (Table 5). Across the whole genome, 34.28% of the genome was identified as a repeat by RepeatMasker. Unplaced contigs and centromere (and centromere-like) regions showed a high coverage of repeat content, close to 100%. Next, telomeres (defined as the region 250kb at the end of each chromosome arm), had a repeat content of approximately 71%, and the non-telomeric/centromeric part of the assembly (containing most of the protein-coding genes) had only 1.4% repeat content.

**Table 5.**
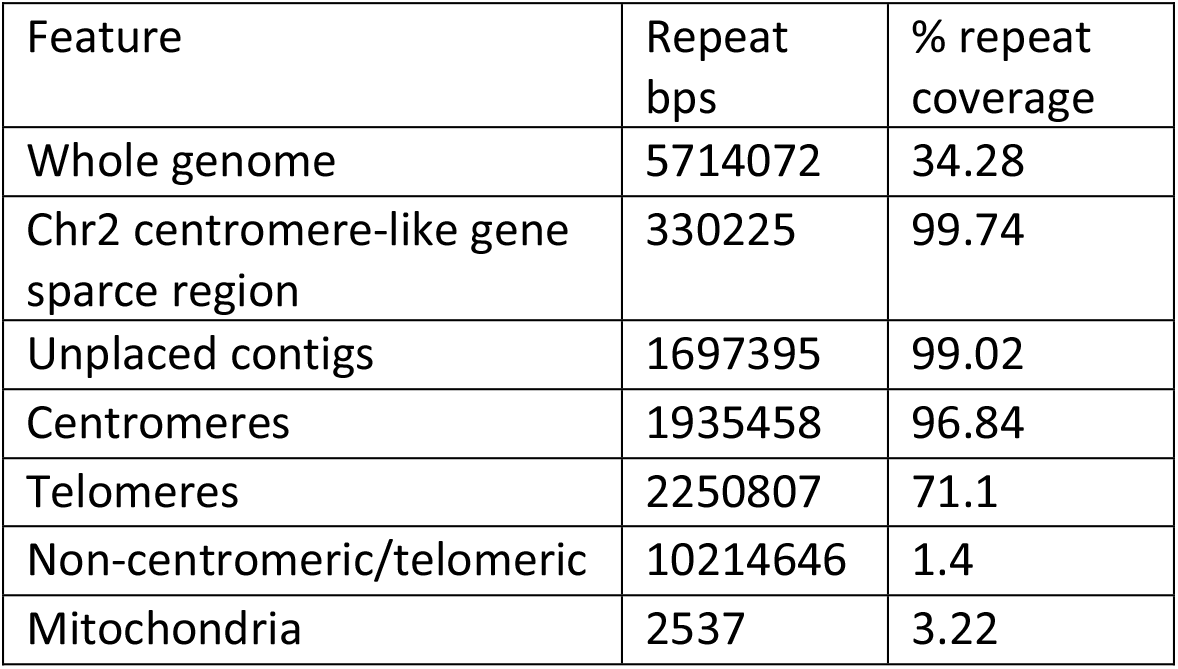
Repeat content for defined regions of the EI 1.0 assembly. Overlapping repeats were merged, so as to not artificially inflate repeat content calculation.

| Feature | Repeat bps | % repeat coverage |
| --- | --- | --- |
| Whole genome | 5714072 | 34.28 |
| Chr2 centromere-like gene sparse region | 330225 | 99.74 |
| Unplaced contigs | 1697395 | 99.02 |
| Centromeres | 1935458 | 96.84 |
| Telomeres | 2250807 | 71.1 |
| Non-centromeric/telomeric | 10214646 | 1.4 |
| Mitochondria | 2537 | 3.22 |

### Enhanced tRNA gene resolution in EI 1.0

We used Liftoff to lift over the NCBI *S. japonicus* annotation (GCF_000149845.2_SJ5) to EI 1.0. The NCBI annotation contained 5215 features classified as ‘genes’, of which 5126 were identified in EI 1.0. Of these 5126 NCBI genes, 4980 were present in only one copy, with 146 of them present in two-copies or more, representing 2897 gene copies. Including all single and multi-copy genes, a total of 7877 NCBI genes were annotated in EI 1.0 (Table 6). tRNAs represented the majority of multi-copy genes. Of the 146 multi-copy genes, only19 of them were protein-coding genes (Supplementary data S2), 118 of them were tRNAs, and 9 were rRNAs. Of the 19 protein-coding genes, 13 of them (representing 73 gene copies) were only partially annotated (lacking 3’ and/or 5’ UTRs). From the 89 genes that were not identified in our assembly, 81 of them were tRNAs, 6 of them were rRNAs, and two were partially annotated coding genes -SJAG_06621 (Tdh1) and SJAG_06597 (Tih1) (Supplementary data S3).

**Table 6.** The number of single- and multi-copy NCBI genes identified by Liftoff in EI 1.0. Multi-copy genes are the number of genes that are present two-times or more, with the number in brackets representing the total number of copies present.

| NCBI genes | All genes | Protein coding | tRNAs | rRNAs |
| --- | --- | --- | --- | --- |
| Only one copy | 4980 | 4857 | 123 | 0 |
| Multi copies | 146 (2897) | 19 (85) | 118 (2623) | 9 (189) |
| Total copies | 7877 | 4942 | 2746 | 189 |

tRNAscan-SE identified 3885 tRNAs *de novo* which overlapped all the 2746 annotated by Liftoff. Of these, 298 were found on Chr1, 2400 on Chr2, 408 on Chr3, 12 on the Mitochondrial genome, and 767 across 3 of the 13 unplaced contigs.

### Other annotation

barrnap and rnammer were used to identify rRNA genes *de novo*. barrnap identified 624 rRNAs, which after merging overlapping loci accounted for 499 non-overlapping loci. rnammer identified 494 rRNAs, all of which overlapped the 499 non-overlapping barrnap loci. From the assembled chromosomes 250 5S rRNA loci were identified on Chr2, 12 18s-28S rRNA loci were identified on Chr3, and no rRNAs were identified on Chr1. Additionally, 237 18S-28S loci were found across 9 of the 13 unplaced contigs.

We used Infernal (v1.1.4) to annotate snRNA and snoRNA genes using previously annotated loci from both *S. japonicus* and *S. pombe*. From this we identified 21 snoRNA loci and 11 snRNA loci, which were distributed evenly across Chromosomes 1 and 2, but only 1 snRNA locus and 2 snoRNA loci were located on Chromosome 3.

### Comparison to JaponicusDB assembly

We used REAPR to map Illumina paired-end reads to both EI 1.0 and JaponicusDB assemblies and looked for assembly errors. REAPR did not produce any breaks across EI 1.0, but produced a total of 10 breaks into the JaponicusDB assembly. Four of these breaks were in small (<200 kb) unplaced scaffolds (supercont5.4 and supercont5.5), two were in Chr I and four were in Chr II (Figure 3). The breaks on Chr I are across the centromere and a tRNA and retro-transposon-rich flanking region, and the four breaks on Chr II were around a 30 kb region containing an array of tRNAs and rRNAs.

**Figure 3.**
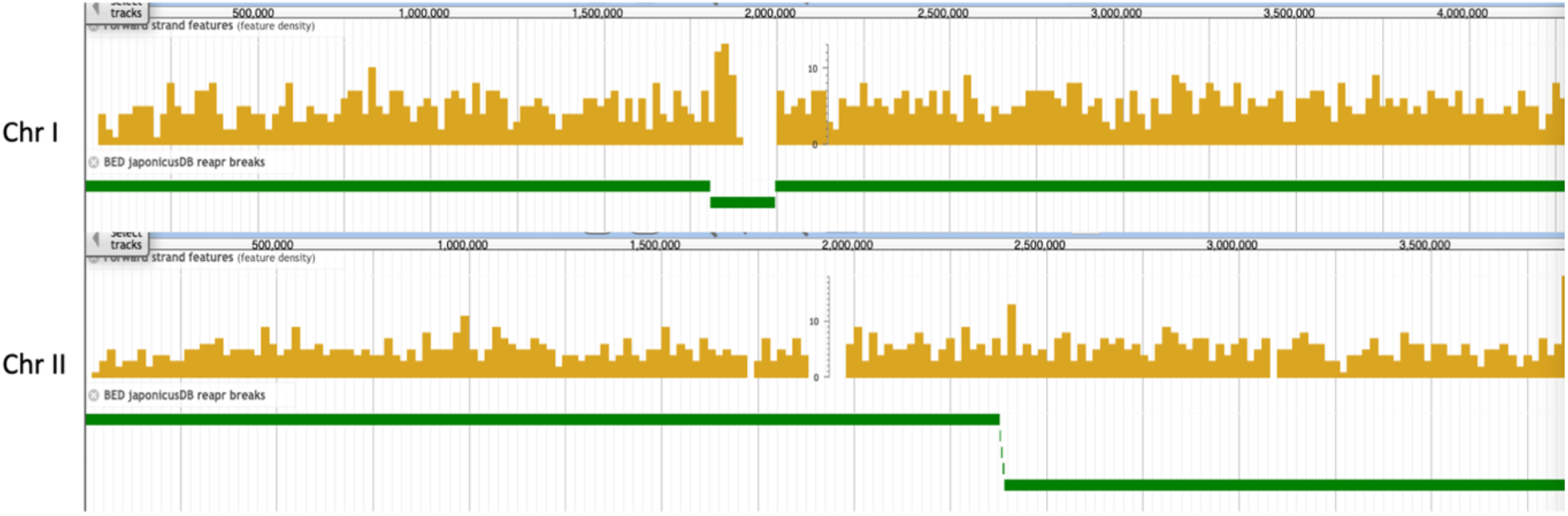
Breaks in the JaponicusDB assembly as suggested by REAPR analyses for Chromosomes I and II. The yellow track is the coverage of genome features in JaponicusDB (coding genes, small RNAs, etc) and the green track represents the contigs produced in the REAPR ‘broken’ assembly. No breaks for identified in Chromosome III.

Considering only the assembled chromosomes, EI 1.0 totalled 16.6 Mb in length, with JaponicusDB chromosomes totalling 11.27 Mb. Using the nucmer alignment (see above), we identified differences in telomere and centromere lengths, along with other large features absent from the JaponicusDB assembly (Figure 1).

### EI 1.0 contains 1.8 Mb of extra telomeric sequence

In total, we resolved 1.84 Mb of extra sequence at the telomeres of all three chromosomes, of which over 1 Mb was attributable to Chromosome 2 (Table 7).

**Table 7.** The estimated length (bps) of additional genomic sequence resolved at the telomeres of EI 1.0 when compared to JaponicusDB. * Denotes the highly-collapsed 5S-rRNA array that might be approximately 2.25 Mb in length.

| Chromosome | 5' arm extension | 3' arm extension | Total |
| --- | --- | --- | --- |
| 1 | 274,257 | 339,994 | 614,251 |
| 2 | 414,184 | 636,974 | 1,051,158 |
| 3 | 162,411 | 14,807* | 177,218 |
| Totals | 850,852 | 991,775 | 1,842,627 |

#### We identified the following features of subtelomeres for each chromosome

Chromosome 1. The 5’ (long arm) telomere was abundant in A-rich repeats, in particularly AACCCT repeats, whilst the 3’ (short-arm) was abundant in G-rich repeats, in particularly GTTTAGG repeats. Reverse complementing these repeats resolves both arms as containing (A)AACCCT repeats.

Chromosome 2. The final 700 kb of the 3’ (long-arm) is rich in LTRs and both telomeres are rich in short-repeats. Also, the final 575 kb is rich in tRNAs. The tRNA arrays generally followed an order of Val-Trp-Thr-Pro-Pro-His-Val-Arg-Ile-His-Ser-Cys-Leu-Pro-Lys, although a shorter version of this (with Val-Arg-Ile-His missing) was also found towards the end of the telomere.

Chromosome 3. The final 45 kb of the 3’ (short-arm) has an array of six 18S-28S rRNA repeat (but see ‘Coverage’ later). Each repeat contains ∼5.9 kb of rRNA genes plus a gap of 2.6 kb between genes. The 5’ (long-arm) starts with an A-rich repeat region of around 60 kb.

### EI 1.0 contains 1.83 Mb of extra centromeric sequence

We identified protein-coding genes with gene standard names, flanking the centromeres in JaponicusDB and then compared the distance between those genes to that in EI 1.0 (Table 8). In total we resolved 1.46 Mb of additional centromeric space than JaponicusDB. Additionally, these centromeric regions in JaponicusDB contained 373,842 Ns, therefore the total increase in resolved chromosomal sequence is 1.83 Mb.

**Table 8.** The distances and differences between the innermost centromere-flanking protein-coding genes of the JaponicusDB assembly and EI 1.0. * -includes 373,842 Ns.

| Chr | Left-flank | Right-flank | JaponicusDB distance | EI 1.0 distance | Difference |
| --- | --- | --- | --- | --- | --- |
| 1 | spt7<br>(SJAG_02927) | rpm1<br>(SJAG_06586) | 230,714 | 610,170 | 379,456 |
| 2 | ppt1<br>(SJAG_06603) | rad31<br>(SJAG_05032) | 160,850 | 644,818 | 483,968 |
| 3 | ser2<br>(SJAG_04449) | rga7<br>(SJAG_01114) | 145,988 | 738,277 | 592,289 |
| Totals |  |  | 537,552* | 1,993,265 | 1,455,713 |

Using our annotation of genomic features, we can see that the centromere of chromosome 2 can be sub-divided into three different regions (Figure 4)

**Figure 4.**
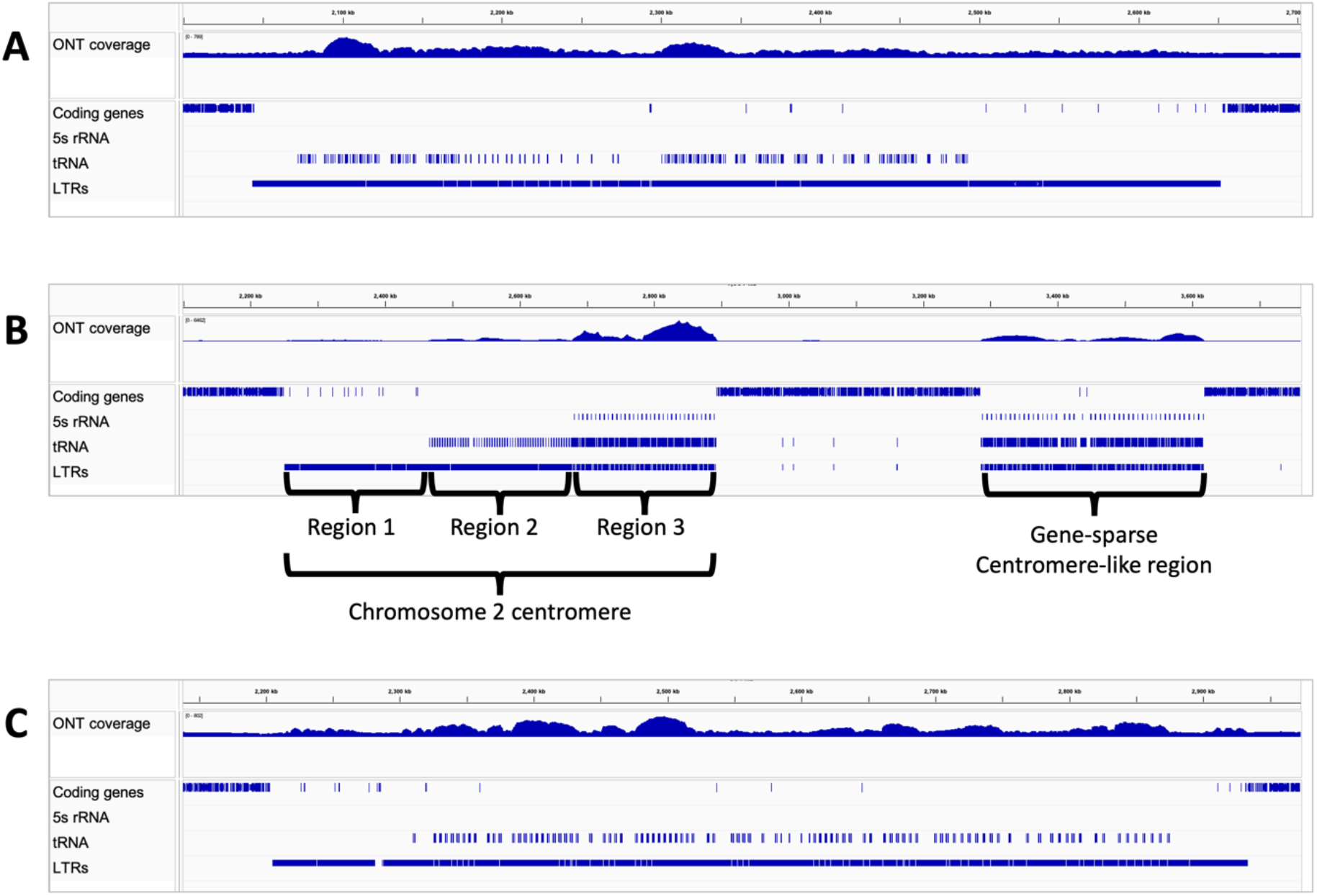
A. ONT long-read coverage (top panel) and gene annotation over centromeres of (A) chromosome 1, (B) chromosome 2 (and gene-sparce centromere-like region), and (C) chromosome 3, showing bordering protein-coding genes, and *de novo* predicted 5S rRNA, tRNA, and LTR loci. In (B) Regions 1, 2, and 3 refer to the three regions defined in the text.

Region 1 -This region of around 215 kb consists of only LTRs, combined with no other repeats or rRNA genes. These LTRs continue across the whole centromere, albeit combined with other centromere-like features.

Region 2 -This region spanning 211 kb, consists of repeated units of approximately 1.2 kb. They contain an Ala-tRNA, a two-exon iMet-tRNA pseudogene, a Gly-tRNA, followed by two A-rich repeats. Each structure is separated by a 2.6 kb gap containing no tRNAs or short-repeats. This region contains LTRs that are also found in region 1 and region 3, but contains no 5S rRNAs.

Region 3 -This part of the centromere (3’), spanning approximately 216 kb consists of 34 copies of a 5.6 kb repeated structure, each containing a T-rich and an A-rich repeat, 9 tRNAs (Tyr-Leu-Glu-Phe-Arg-Lys-Gly-Ile-Gln), an array of three 5S rRNAs, followed by another array of 9 tRNAs (Glu-Leu-Asn-Met-Thr-Asp-Ser-Val-Ala) (Figure 4).

tRNA arrays on Chromosome 3 differed from Chromosome 2 by being much shorter, occurring in an array of five re-occurring tRNAs (Gln-Pro-Thr-Ser-iMet(pseudogene)-Gln). tRNA arrays on Chromosome 1 tended to occur in 3kb repeated tRNA structures (Ala-iMet-Ala-Arg-Pro-Gly), although these structures tended to be reversed at the 3’ end of the centromere.

### Chromosome 2 contains a centromere-like gene sparse region

We located a region of approximately 331 kb on Chromosome 2 that had not been resolved in previous *S. japonicus* assemblies. This region is rich in both tRNAs and 5S rRNAs and only contains one protein-coding gene (in two copies) -SJAG_06608, annotated as a chromo/chromo shadow domain family gene. This part of the chromosome consisting of 46 repeated 6.4 kb tRNA/rRNA structures almost identical to that found at the 3’ end of the centromere (Region 3 above) and differed only in that they tended to be longer (e.g., 6.84 kb, compared to 5.64 in the centromere), and the two short repeats were both T-rich and were further apart than in the centromere (Figure 5.).

**Figure 5.**
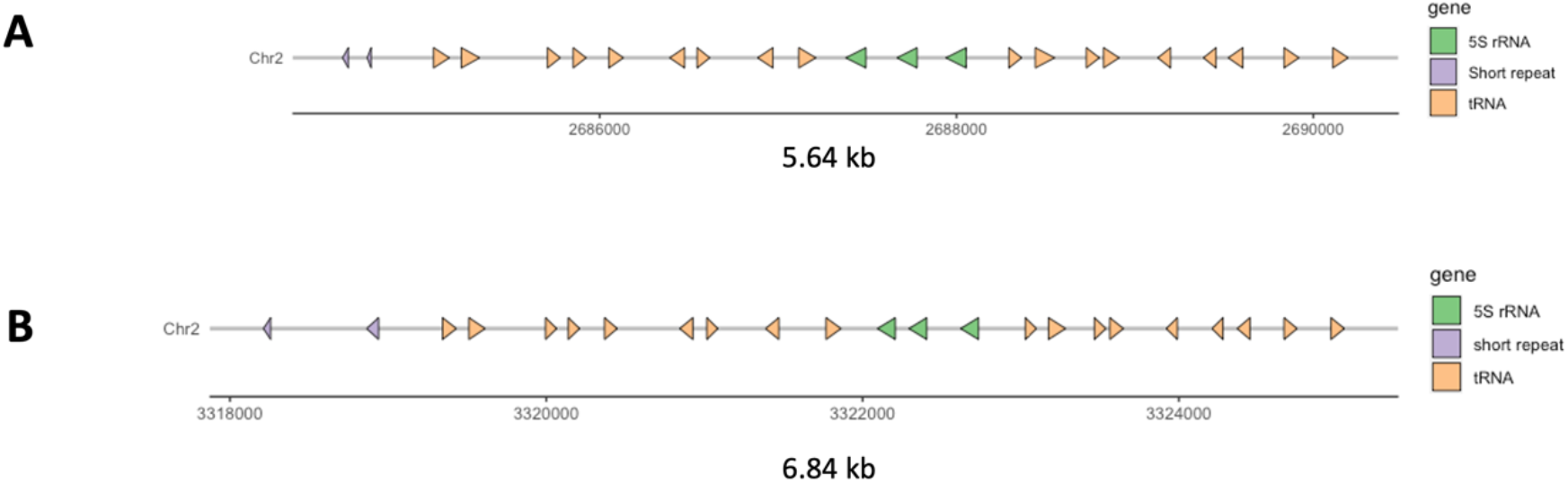
Typical structure of a tRNA arrays from Chromosome 2 in (A) the centromere region 3, and (B) the centromere-like gene sparse region. The array starts with two short repeats, followed by 9 tRNAs, three 5S rRNAs, and ends with nine more tRNAs.

### Coverage analyses suggests collapsed tRNA and 5S rRNA arrays

We calculated long-read coverage across all three chromosomes along with that for only the coding region (Table 9) and compared it to that of other regions of interest on chromosome 2 (Figure 4, Table 10).

**Table 9.** Mean per-base coverage and standard deviation (SD) of ONT long-reads across whole chromosomes and coding regions only of EI 1.0. Coding regions are defined by the start position of the first and end position of the last BUSCO ortholog on each chromosome arm.

| Chr | Whole chromosome (SD) | Coding regions (SD) |
| --- | --- | --- |
| 1 | 161 (40) | 176 (15) |
| 2 | 172 (94) | 176 (17) |
| 3 | 168 (78) | 173 (16) |

**Table 10.** Mean per-base coverage and standard deviation (SD) of ONT long-reads and across centromere features and centromeric-like regions on chromosome 2.

| Interval (length) | ONT coverage (SD) | Comment |
| --- | --- | --- |
| Chr2:2250038-2891897 (642 kb) | 248 (240) | Chr2 centromere |
| Chr2:2250038-2464999 (215 kb) | 151 (16) | Region 1 – Chr 2 LTR region |
| Chr2: 2465000-2675532 (211 kb) | 111 (16) | Region 2 – Chr 2 centromere tRNA array |
| Chr2:2675533-2891897 (216 kb) | 479 (301) | Region 3 – Chr 2 centromere tRNA and rRNA arrays |
| Chr2:3285699-3617048 (331 kb) | 202 (94) | Chr2 centromere-like gene sparse region |

Mean long-read coverage across the three chromosomes varied between 161x to 172x, with standard deviations (SD) of between 40 and 94. When only considering coding regions, the coverage was much more even. Mean coverage for coding regions varied between 173x to 176x, with SD of between 15 and 17. The mean coverage across only the coding regions of Chromosome 2 was 176 (SD 17), whilst that of the centromere was 248 (SD 240), suggesting large variation in coverage between coding and non-coding regions. We split the centromere into three regions, paying particular attention to regions 2 and 3 (Table 10). Region 2 had coverage slightly below both the centromere mean and whole chromosome mean, and Region 3 which also contained tRNAs and a large array of rRNAs, had coverage 2.7 times that of the chromosome mean (Table 10 and Figure 4). We also examined coverage over the 331 kb centromere-like gene sparse region, which had coverage slightly above the mean, albeit with a high SD. Taking coverage into consideration, Region 2 might have a slightly shorter length of 135 kb and Region 3 might have a greater length of 601 kb, increasing the estimated centromere size from 642 kb to 952 kb.

### Chromosome 3 telomeric rRNA array is likely not fully resolved

Additionally, we also looked at 18S-28S rRNA arrays in the assembly. These were present in a 49 kb region at the 5’ end of Chromosome 3 and the 71kb unplaced contig tig00000022. tig00000022 is a reverse complement assembly duplication of the 3’ end of Chromosome 3 and together likely represent a collapsed 18S-28S rRNA array that was not merged during assembly. Additionally, 8 other unplaced contigs contained only 18S-28S rRNA arrays (Table 11). The cumulative length of the assembled 18S-28S rRNAs is 1.05 Mb. Assuming that all these sequences represent a single telomeric 18S-28S rRNA array from Chromosome 3 and accounting for coverage and length of these sequences, it is likely that the actual size of this rRNA array is approximately 2.25 Mb in length (Supplementary Table S4).

**Table 11.** Mean per-base coverage and standard deviation (SD) of ONT long-reads across 18S-28S rRNA arrays.

| Interval | Mean coverage (SD) | Comment |
| --- | --- | --- |
| Chr3:3749417-3796598 (47 kb) | 725 (337) | Chromosome 3 telomeric 18S-28S rRNA array |
| Unplaced 18S-28S rRNA arrays (1 Mb) | 377 (398) | 9 unplaced contigs with 18S-28S rRNA arrays across most of sequence. |

## Discussion

### Genome size

We have sequenced, assembled, and annotated a *de novo* genome assembly for *Schizosaccharomyces japonicus* (EI 1.0), which is more complete and contiguous than previous versions. Each chromosome is represented by a single telomere-to-telomere contig. The chromosomes (totalling 14.8 Mb) contain roughly 3.6 Mb (30.5%) more sequence than the chromosomes in the current reference sequence (JaponicusDB SJ5, size 11.2 Mb). This additional sequence consists of 1.84 Mb more sub-telomeric sequence, 1.45 Mb more centromeric sequence and a 331 kb centromere-like region on chromosome 2 that had not been previously described. The assembly has 1.97 Mb more repeat content than JaponicusDB, therefore >50% of the size increase is down to better resolution of repeat content or repeat-rich regions (i.e., repeats, telomeres, and centromeres), rather than gene content (EI 1.0 contains only 0.7% more single-copy orthologs than JaponicusDB). When accounting for coverage over centromeres and telomeres, the true genome size might be closer to 18.12 Mb (Supplementary Table S4).

### Chromosome 2 centromere-like gene sparse region

The three assembled chromosomes in the JaponicusDB assembly were inferred from genetic linkage within the seven largest supercontigs, with the inferred centromeres linked by stretches of 120 kb of ‘N’s. The centromere gaps can be seen in the genome alignments for all three chromosomes (Figure 1). The final two supercontigs to be closed on the JaponicusDB assembly were on chromosome 2 (supercontigs 7A and 5D), which corresponds to the 331 kb gene-sparse region we identify on chromosome 2. This is ∼390 kb downstream from the centromere and both features are shown as gaps in the genome alignments (Rhind 2011, Supp Figure 2, and Figure 1). Further evidence of this region not being properly assembled in JaponicusDB is shown by the REAPR analysis which shows four misassembly breaks across this region (Figure 3). This region displays most resemblance to the 3’ end of the centromere on the same chromosome, characterised by being rich in both 5S rRNAs and tRNAs across its full length, of which the tRNA array order is identical to that of the centromere. However, the composition of and the distance between the short repeats differs from that of the centromere, providing confidence that this region is distinct and correctly assembled. The centromeres of chromosomes 1 and 3 are rich in tRNAs and LTRs but have no rRNA arrays, suggesting that the centromere-like gene sparce region on chromosome 2 originated from the centromere of chromosome 2. Long-read coverage analyses of the region suggests that the actual length may be 1.3x longer, around 430 kb. Liftoff annotated 43 copies of the rRNA gene SJAG_16156 across this region. SJAG_16156 can be located on JaponicusDB Chromosome II and is part of a 13 kb rRNA/tRNA array (containing 3 rRNAs and 7 tRNAs) flanked by genes SJAG_04837 and SJAG_04436. These two genes also flank the 331 kb centromere-like gene sparce region, providing evidence of the collapsing of rRNA and tRNA arrays in JaponicusDB.

### tRNA arrays

Long arrays of tRNAs are found across the majority of each centromere and the centromere-like gene sparse region on Chromosome 2. The exceptions to this are in the centromere of Chromosome 1, where they are absent from the last 160 kb, and the centromere of Chromosome 2, where they are absent from the first 215 kb (Figure 4). There is also a 575 kb tRNA array on the 3’ telomere of Chromosome 2. Additionally, tRNAs are also present on unplaced contigs tig00000004, tig00000005, and tig00000029. Compared to previous assemblies, EI 1.0 has 2623 extra annotated copies of tRNA genes, comprising 90% of all novel annotated genes in this assembly. It had been proposed that S. japonicus contained no tRNAs with Ala(AGC) anticodon, which is the most abundant in other yeasts, and decoded the GCU Ala codon exclusively through a species-specific wobble Ala(TGC), and Ala(CGC) (Iben & Maraia, 2012). We indeed found Ala(TGC) (106 copies across all three chromosomes) and Ala(CGC) tRNAs (47 copies, all on Chromosome 1). However, we also discovered 80 copies of Ala(AGC), all situated on Chromosome 2, within the centromere and the centromere-like gene sparse region. Both of these regions are poorly resolved in the previous JaponicusDB SJ5 assembly, providing further evidence of the advantages of long-read sequencing assembly.

### rRNA arrays

EI 1.0 contains three rRNA arrays -two 5S rRNA arrays on Chromosome 2 (one in the centromere, and one in the centromere-like gene sparse region), and one 18S-28S rRNA array on the 3’ (short-arm) of Chromosome 3. rRNAs are absent from Chromosome 1. The 5S rRNAs in the centromere of Chromosome 2 are restricted to a 216 kb tandem array at the 3’ end, which is represented by 34 repeated modules of three 5S rRNAs (with one exception, where only one 5S rRNA is present). The centromere-like gene sparse region on Chromosome 2 has a 5S rRNA tandem array made up of 46 repeated modules of between one and four rRNAs in each module. Modules with three rRNAs dominated the first 116 kb of this region, with modules of four rRNAs dominating the final 215 kb. In all but three modules, the first rRNA shows a high degree of homology to JaponicusDB gene SJAG_16156. In each of the three exceptions, the rRNA module contained only one or two rRNAs. The remaining rRNAs in each module did not have homologous genes in JaponicusDB (defined by the lack of Liftoff annotations over each one), suggesting a lack of rRNA array resolution in that assembly. Chromosome 3 has a 45 kb array of 18S-28S rRNAs at the very end of the short-arm telomere. The array is formed of six modules of reoccurring 18S-28S pairs. This is the only place in the assembled chromosomes that has this pattern of rRNAs, but it should be noted that 9 of the 13 unplaced contigs also have the same pattern of 18S-28S pairs across their full length (including tig00000022 which is a reverse complement of the 3’ end of chromosome 3). The cumulative length of these contigs is ∼1 Mb (2.25 Mb when adjusted for coverage) and likely represent un-scaffolded regions of the Chromosome 3 sub-telomere. This is consistent with the physically marked position of the 18S – 28S rRNA array in *S. japonicus* (Yam et al., 2013).

We have completed the first telomere-to-telomere genome assembly of the fission yeast *S. japonicus* and described a hitherto undescribed genome architecture. Previously unseen features include vastly expanded repeat arrays, including those encoding tRNAs and rRNAs, compared to other fission yeast species. Another notable feature is the centromere-like gene-sparce region on the long arm of chromosome 2, with as yet unknown implications for the evolution of centromeres and perhaps chromosome number changes. It will be interesting to explore the mechanisms and evolutionary pressures that have led to and fixed this huge increase in repeat number, as well as how they contribute to the physiology and life-style of this unique model species.

This telomere-to-telomere genome assembly required the resolving of complex repetitive features (including tRNA and rRNA arrays) which was possible due to the availability of long ONT sequencing reads (>50 kb). However, even with such long reads, coverage analysis demonstrates and quantifies some collapse of the highly repetitive regions (Figure 4, Table 10, Table 11). It is worth noting that there is likely to be diversity in repeat copy number between *S. japonicus* isolates and even within a culture. Understanding these differences will be aided be increases in nanopore sequencing read length and accuracy. Nanopore sequencing generates read lengths that reflect the length of purified DNA fragments with (theoretically) no limit on read length. Therefore, in the future, nanopore sequencing datasets with even longer reads will allow increasingly accurate genome assemblies and shed more light on as-yet undiscovered genomic diversity.

## Acknowledgements

The authors would like to thank Darren Heavens (Earlham Institute) for assistance during nanopore library preparation and sequencing.

GE assembled and annotated the assembly and carried out all bioinformatic analyses. PSW created the cell culture and carried out HMW DNA extraction. CN, SO, and FU assisted with experimental design. GE, CN, SO, and FU contributed to writing and reviewing the manuscript.

## Data availability

All new data for the assembly can be located under ENA Project PRJEB63404. The genome assembly can be found under accession number CATPIX010000000.

## Supplementary Data

Supplementary data S1(Data_S1_s_japonicus_gff.tgz) – Genome annotation files for *S. japonicus* EI 1.0

Supplementary data S2 – Multi-copy protein coding genes found in EI 1.0.

Supplementary data S3 – Unmapped genes. A list 89 genes from the NCBI SJ5 annotation, not found in EI 1.0.

Supplementary table S4 – Calculations of lengths and coverage-adjusted feature lengths in all contigs, plus the Chromosome 2 centromere and the Chromosome 3 18S-28S rRNA array.

